# Web-based design and analysis tools for CRISPR base editing

**DOI:** 10.1101/373944

**Authors:** Gue-Ho Hwang, Jeongbin Park, Kayeong Lim, Sunghyun Kim, Jihyeon Yu, Sang-Tae Kim, Roland Eils, Jin-Soo Kim, Sangsu Bae

## Abstract

**Background:** As a result of its simplicity and high efficiency, the CRISPR-Cas system has been widely used as a genome editing tool. Recently, CRISPR base editors, which consist of deactivated Cas9 (dCas9) or Cas9 nickase (nCas9) linked with a cytidine or a guanine deaminase, have been developed. Base editing tools will be very useful for gene correction because they can produce highly specific DNA substitutions without the introduction of any donor DNA, but dedicated web-based tools to facilitate the use of such tools have not yet been developed.

**Results:** We present two web tools for base editors, named BE-Designer and BE-Analyzer. BE-Designer provides all possible base editor target sequences in a given input DNA sequence with useful information including potential off-target sites. BE-Analyzer, a tool for assessing base editing outcomes from next generation sequencing (NGS) data, provides information about mutations in a table and interactive graphs. Furthermore, because the tool runs client-side, large amounts of targeted deep sequencing data (>100MB) do not need to be uploaded to a server, substantially reducing running time and increasing data security. BE-Designer and BE-Analyzer can be freely accessed at http://www.rgenome.net/bedesigner/ and http://www.rgenome.net/be-analyzer/respectively

**Conclusion:** We develop two useful web tools to design target sequence (BE-Designer) and to analyze NGS data from experimental results (BE-Analyzer) for CRISPR base editors.

## Background

CRISPR-Cas (clustered regularly interspaced short palindromic repeats and CRISPR associated), an immune system in bacteria and archaea that targets nucleic acids of viruses and plasmids, is now widely used as a genome editing tool because of its convenience and high [1–3]. The most popular endonuclease, type II CRISPR-Cas9, makes DNA double-stranded breaks (DSBs) at a desired site with the help of its single-guide RNA (sgRNA) [4,5]. The DSBs provoke the cell’s own repair systems: error-prone non-homologous end joining (NHEJ) and error-free homology-directed repair (HDR), resulting in gene knock-out and knock-in (or gene correction), respectively. However, it is relatively difficult to induce gene corrections such as one nucleotide substitutions because HDR occurs rarely in mammalian cells compared to NHEJ [6]. Furthermore, Cas9 can frequently induce DSBs at undesired sites with sequences similar to that of the sgRNA [7,8].

Recently, CRISPR-mediated base editing tools have been developed. These tools enable the direct conversion of one nucleotide to another without producing DSBs in the target sequence and without the introduction of donor DNA templates. The initial base editors (named BEs), composed of dCas9 [9] or nCas9 [10] linked to a cytidine deaminase such as APOBEC1 (apolipoprotein B editing complex 1) [11] or AID (activation-induced deaminase) [12], substitute C for T. Later, adenine base editors (ABEs) were constructed by using tRNA adenine deaminase (TadA), evolved to enable the direct conversion of A to G in DNA [13]. Because of their ability to make highly specific DNA substitutions, these base editing tools will be very useful for gene correction [14–17], but to the best of our knowledge, a user-friendly web-based tool for their design and analysis has not yet been developed.

Here, we present dedicated web toolkits, named BE-Designer and BE-Analyzer, to aid researchers in choosing sgRNAs to target desired DNA sequences and to assess base editing outcomes from next generation sequencing (NGS) data. BE-Designer provides researchers with a list of all possible sgRNAs for targeting given input DNA sequences, along with useful information including their potential off-target sites, for 250 registered organisms, presently. After introducing CRISPR base editors into a population of cells, researchers ultimately perform targeted deep sequencing to measure mutation efficiencies and analyze DNA mutation patterns. BE-Analyzer analyzes and summarizes NGS data in a user’s web browser; because of the advantages of JavaScript, there is no need to upload data to a server or install local tools. BE-Analyzer also optionally accepts control data from CRISPR-untreated cells and displays the output in an additional nucleotide mutation table so that users can easily compare the data from CRISPR-treated and untreated cells.

## Implementation & Results

### BE-Designer

BE-Designer is a sgRNA designing tool for CRISPR base editors. BE-Designer rapidly provides a list of all possible sgRNA sequences from a given input DNA sequence along with useful information: possible editable sequences in a target window, relative target positions, GC content, and potential off-target sites. Basically, the interface of BE-Designer was developed using Django as a backend program.

### Input panels in BE-Designer

BE-Designer presently provides analysis for CRISPR base editors based on SpCas9 from *Streptococcus pyogenes*, which recognizes 5’-NGG-3’ protospacer-adjacent motif (PAM) sequences, as well as SpCas9 variants: SpCas9-VQR (5’-NGAN-3’), SpCas9-EQR (5’-NGAG-3’), SpCas9-VRER (5’-NGCG-3’) [18]. BE-Designer also provides analysis for CRISPR base editors based on SaCas9 from *Staphylococcus aureus* (5’-NNGRRT-’3) and its engineered form, SaCas9-KKH (5’-NNNRRT-’3) [19]. Currently, BE-Designer supports sgRNA design in 255 different organisms, including vertebrates, insects, plants, and bacteria. Users can input DNA sequences directly in the target sequence panel of the web site or upload a text file containing DNA sequences. The DNA sequence should be a raw string comprised of IUPAC nucleotide codes or FASTA formatted text. By using an analysis parameter, users can manually select the type of base editor, either BE or ABE, and the base editing window in the target DNA (Fig. 1A).

**Fig 1.**
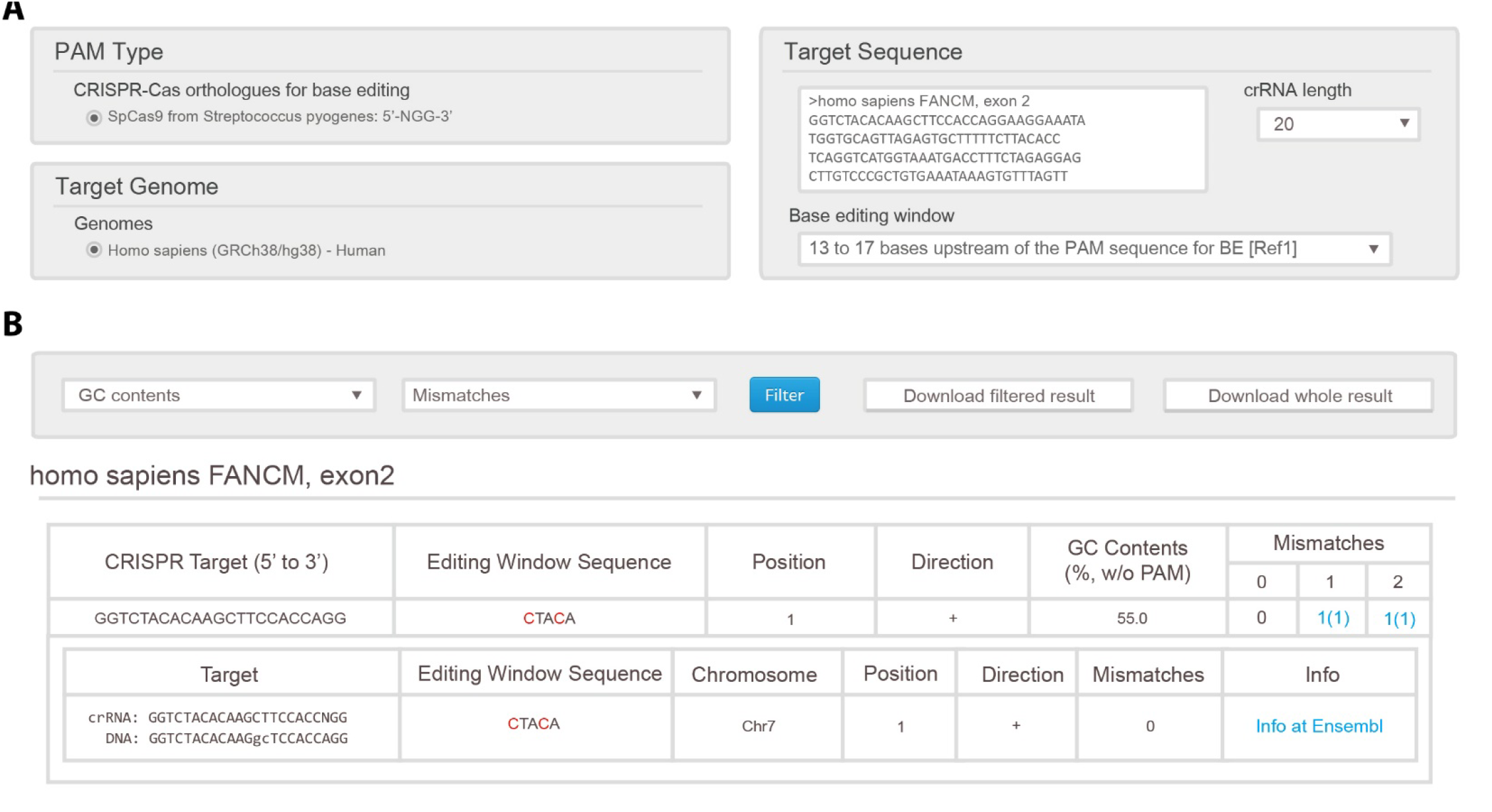
Scheme of BE-Designer. **a** In the BE-Desinger, users collect PAM type, genome of organism and base editing window and input the sequence that is found to target sequence. And, after users click the submit button, BE-Designer display results page. **b** The results page presents the possible target sequences and thoses information that contain expected sequence in eiditing window sequence, position, direction, GC contents and the count of off-target at up to 2 mismatches. Clicking the number of off-taget, users can receive the off-target’s information. And if the organism’s genome is in the Ensemble, BE-Designer connects to the Ensemble page.

### Selection of sgRNAs

Within a given DNA sequence, BE-Designer finds all possible target sites based on input parameters; in the base editing window, target nucleotides are highlighted in red, and their relative position and GC content are indicated. BE-Designer then invokes Cas-OFFinder [20] to search throughout the entire genome of interest for possible off-target sequences that differ by up to 2 nucleotides from the on-target sequences.

### Result visualization

BE-Designer produces a result table that contains the target sequences with useful information (Fig. 1B). BE-Designer uses AJAX (Asynchronous JavaScript and Extensible Markup Language) to show results instantly; thus, users can filter the results according to GC content and mismatch numbers without refreshing the whole web page. In addition, if the Ensembl annotation is available for the given reference genome, BE-Designer offers a link to the corresponding Ensembl genome browser web page that displays the sequence information near any off-target loci.

### BE-Analyzer

Due to its high sensitivity and precision, targeted deep sequencing is the best method for assessing the results of base editing. BE-Analyzer accepts targeted deep-sequencing data and analyzes them to calculate base conversion ratios. In addition to the interactive table and graphs showing the results, BE-Analyzer also provides a full list of all query sequences aligned to a given wild-type (WT) sequence, so that users can confirm mutation patterns manually. The BE-Analyzer interface was also developed using Django as a backend program. The core algorithm of BE-Analyzer was written in C++ and then transcompiled to asm.js with Emscripten (http://kripken.github.io/emscripten-site/)

### Input panels in BE-Analyzer

NGS data are typically composed of a pair of Fastq files from paired-end sequencing, or a single Fastq file from single-read sequencing. BE-Analyzer allows both types; if the input is a pair of Fastq files, BE-Analyzer first merges them by the JavaScript port of fastq-join, a part of ea-utils (https://code.google.com/archive/p/ea-utils/). As an option, users can additionally upload data from a CRISPR-untreated control to compare it with data from the treated sample (Fig. 2A). In this case, BE-Analyzer analyzes the two datasets simultaneously and compares them to exclude background mutations found in the control sample.

**Fig 2.**
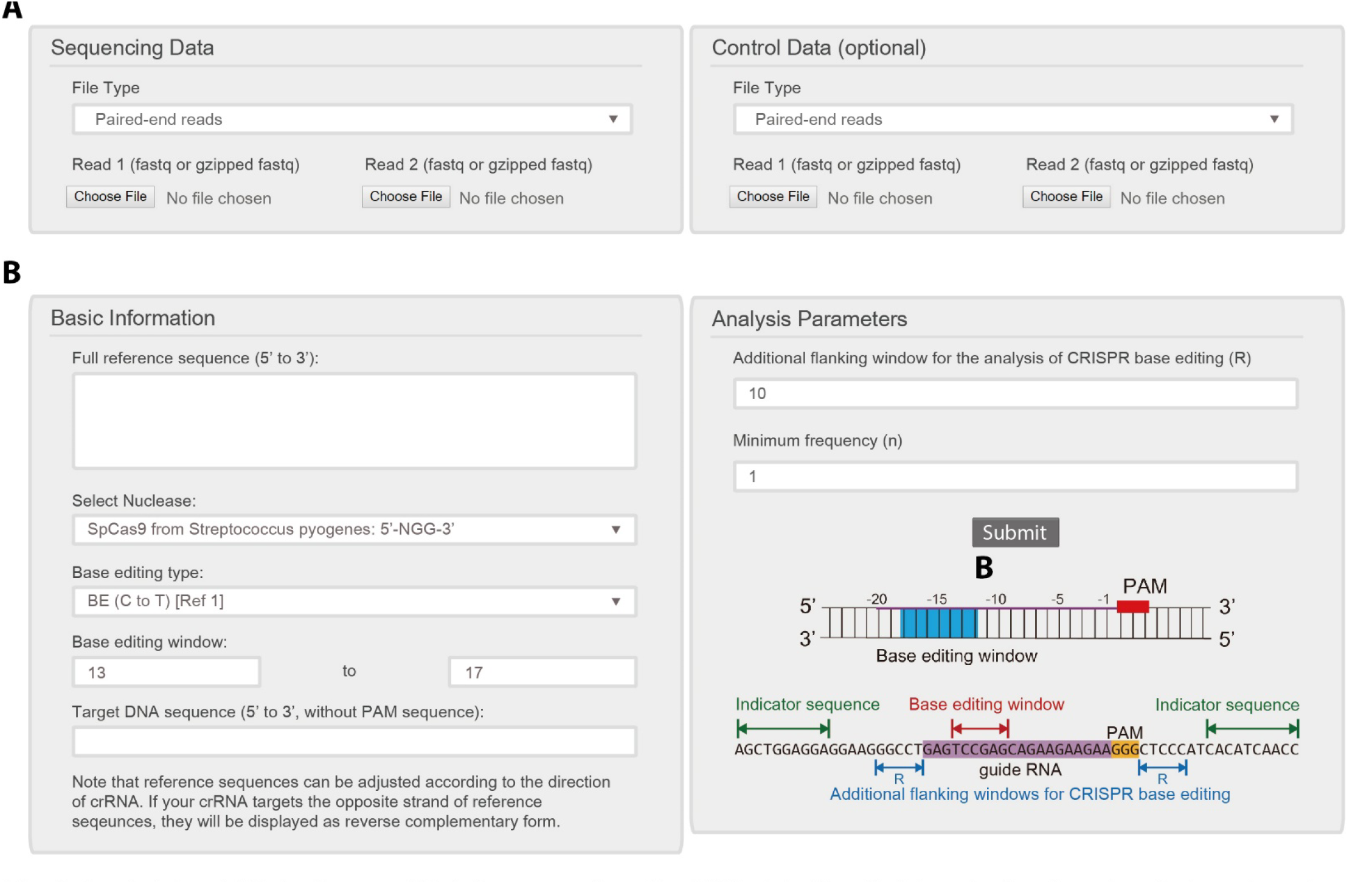
Input data of BE-Analyzer. **a** BE-Anlayzer receives the NGS data files that is paired-end reads, single-end reads, or fastq-joined file. **b** The basic information about the sequence data like reference sequence, nuclease type, base editing type and target seqeucne is received. The Additio nal flanking window is an additional dispalyed range at both end of target sequence in results. BE-Analyzer filter sequences of which reads is lower than Minimum frequence.

To analyze query sequences in NGS data, BE-Analyzer requires basic information: a full WT sequence for reference, the type of base editor, the desired base editing window, and the target DNA sequence (Fig. 2B). Previous studies have reported the optimal target window for each base editor. For example, BE3 usually induces base conversion in a region ranging from 13 to 17 nucleotide (nt) upstream of the PAM, and TARGET-AID is most efficient within a region 15 to 19 nt upstream of the PAM. Basically, BE-Analyzer provides the optimal default values with reference to previous studies, but users can freely revise the value manually. On the other hand, it has been reported that base editors can introduce substitutions outside of the DNA target sequences at a low frequency [12]. Therefore, BE-Analyzer is implemented to allow additional flanking windows on each side of the target for analysis by the use of a relevant parameter.

### Analysis of NGS data

From uploaded NGS data, BE-Analyzer first defines 15-nt indicator sequences on both sides of the given reference sequence; only identified queries that have both indicator sequences, with ≤1 nt mismatches, are collected. Then, BE-Analyzer counts the recurrent frequency of each sequence and sorts queries in descending order. In this procedure, sequences with frequencies below the minimum are discarded. Each sequence is aligned to the reference sequence with EMBOSS needle (https://www.ebi.ac.uk/Tools/psa/emboss_needle/). As a result, the aligned sequences are classified into four different groups based on the presence of a hyphen (-). If hyphens are found in the reference sequence or query, the query is classified as an insertion or deletion by a comparison of the number of hyphens in the two sequences. If hyphens (inserted or deleted sequences) are not found in a given target window including the additional flanking regions, the query is referred as a WT sequence. Otherwise, the queries that contain a few mismatched nucleotides in the given target window are classified as substitutions.

Among the query sequences defined as substitutions, if there are desired base conversions, i.e. C to D (A, G, or T) for BE and A to G for ABE, in the given target window, BE-Analyzer further analyzes them to calculate the ultimate base editing efficiency and to display the base editing patterns in interactive tables and graphs. A table showing statistics, base editing efficiencies, information about expected amino acids, and the categorized align result tab are displayed using Bootstrap library. Bar graphs and heat maps of substitution patterns are visualized using Plotly.js (https://plot.ly/javascript/).

### Result visualization

The results are summarized as a table with 9 columns (Fig. 3A): (i) ‘Total Sequence’ indicates the number of all reads present in the Fastq file, (ii) ‘With both indicator sequences’ indicates the number of reads having both indicator sequences, (iii) ‘More than minimum frequency’ indicates the number of reads that remain after the reads that appear with less than the minimum frequency are removed, (iv, v, vi) ‘Wild type’, ‘Insertions’, and ‘Deletions’ indicate the number of reads in each category, (vii) the 7th column indicates the number of reads having at least one base substitution, (viii) the 8th column indicates the number of reads that have nucleotide conversions induced by CRISPR base editors in target windows, and (ix) the 9th column indicates the intended substitution rate (such as ‘C to T Substitution Rate’), obtained by dividing the number of reads that have intended conversions in the base editing window with the number of reads above the minimum frequency (3rd column).

**Fig 3.**
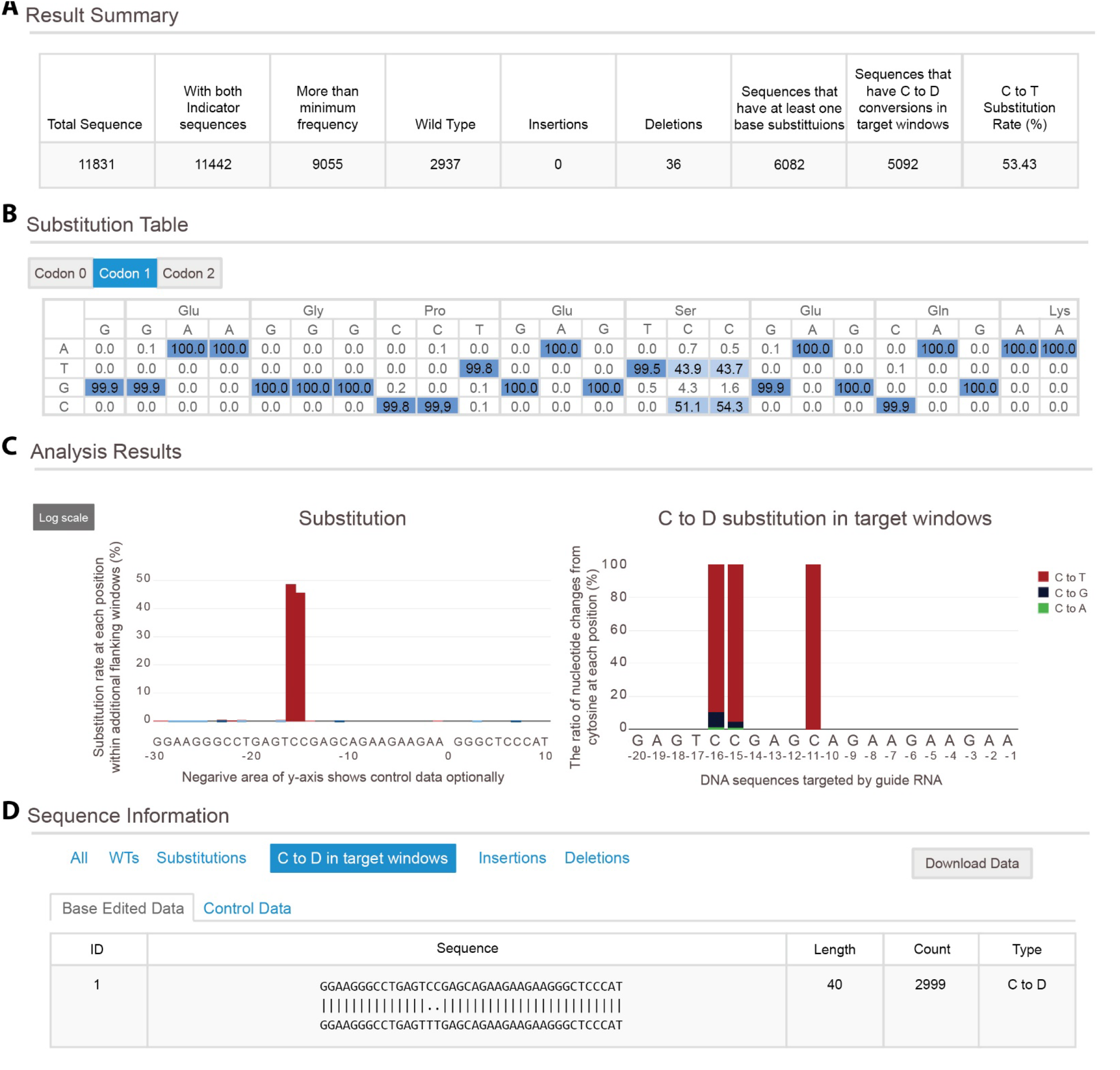
Results of BE-Analyzer about BE3. **a** The table includes the file sequence count and mutation count **b** Substitution table shows the nucleotide mutation at each position and the expected amino acid sequence. The buttons control the start of amino acid sequence. **c** Substituion rate at each position is displayed in substitution graph in additional flanking sequence and Substituion pattern (C to D) is displayed in C to D substitution graph in target sequence. And BE-Analyzer also displays Substitution pattern between nucleotide. **d** All filtered sequences are aligned with reference sequence.

For base editing, it is crucial to know how the mutation of one or a few nucleotides changes the amino acid sequence. To address this issue, BE-Analyzer provides the expected amino acid sequences for three different reading frames, so that users can select among three possible start positions (Fig. 3B). For each nucleotide, BE-Analyzer displays the nucleotide mutation rate in detail, highlighted with a color gradient.

Although cytidine deaminases mainly introduce C to T transitions in the base editing window, C to A or G transitions may also occur in flanking regions with low probability. Thus, BE-Analyzer shows the substitution rate at each site in the flanking windows and the C to D transition pattern in the target windows (Fig. 3C). In the C to D substitution graph, each transition pattern is presented with its percentile rate, and the type of transition indicated by color (red-black-green). Optionally, if users previously uploaded data from a CRISPR-untreated control, BE-Analyzer displays the substitution rate at each of those sites in the negative direction. Furthermore, for users’ convenience, BE-Analyzer shows substitution patterns within the flanking windows with a heat map, which enables visualization of the dominant substitution patterns as well as background patterns.

At the bottom of the results page, a list of categorized sequence reads aligned to the reference sequence is presented (Fig. 3D). Users can confirm all filtered sequences from the input data in this table and can also save the results by clicking the ‘Download Data’ button.

## Conclusions

BE-Designer is an easy-to-use web tool for optimal selection of sgRNAs in a given target sequence. It identifies all possible target sequences in a given sequence and displays information about each target sequence, including predicted mutation patterns, mutation positions, and potential off-target sites. Users can easily select the optimal sgRNA sequence for current base editors. BE-Analyzer is another web tool for instant assessment of deep sequencing data obtained after treatment with base editors. BE-Analyzer instantly analyzes deep sequencing data at a client-side web browser and displays the results using interactive tables and graphs for users’ convenience. Useful information, including the ratio of intended conversions, transition patterns, and sequence alignments, is provided so that users can easily infer how frequently and where intended or unwanted substitutive mutations are generated.

## Abbreviations

CRISPR-Cas: clustered regularly interspaced short palindromic repeats and CRISPR associated
DSB: DNA double-stranded breaks
sgRNA: single-guide RNA
NHEJ: non-homologous end joining
HDR: homology-directed repair
ABEs: adenine base editors
TadA: tRNA adenine deaminase
NGS: next generation sequencing
PAM: protospacer-adjacent motif
WT: wild-type

## Declarations

### Acknowledgements

We thank Dr. M. Schlesner at DKFZ for helpful discussion.

### Availability of data and materials

Example NGS data is freely accessible from the web site (http://www.rgenome.net/be-analyzer/example.

### Authors’ contributions

G.-H.H., J.P. and S.B. designed this project. G.-H.H. and J.P. constructed the web tools reported in this study. K.L., S.K., J.Y. and S.-T.K. gave critical comments on the web panel. R.E., J.-S.K. and S.B. supervised the research. G.-H.H., J.P. and S.B. wrote the manuscript with the help of others.

### Ethics approval and consent to participate

Not applicable.

### Consent for publication

Not applicable.

### Additional files

Additional file 1: Figure S1. The internal programs used for implementation of BE-Designer and BE-Analyzer.

Additional file 2: Figure S2. The workflow for classifying query sequences in BE-Analyzer.

